# Seasonal variation in brain mu-opioid receptor availability

**DOI:** 10.1101/2020.05.20.104349

**Authors:** Lihua Sun, Jing Tang, Heidi Liljenbäck, Aake Honkaniemi, Jenni Virta, Janne Isojärvi, Tomi Karjalainen, Tatu Kantonen, Pirjo Nuutila, Jarmo Hietala, Valtteri Kaasinen, Kari Kalliokoski, Jussi Hirvonen, Harry Scheinin, Semi Helin, Kim Eerola, Eriika Savontaus, Emrah Yatkin, Juha O. Rinne, Anne Roivainen, Lauri Nummenmaa

## Abstract

Seasonal rhythms influence mood and sociability. The brain μ-opioid receptor (MOR) system modulates a multitude of seasonally varying socioemotional functions, but its seasonal variation remains elusive with no previously reported *in vivo* evidence. Here, we studied the seasonal effects on brain MOR availability via analysing a dataset (n=204) of [^11^C]carfentanil positron emission tomography (PET) scans of healthy volunteers. We found that seasonally varying daylength had an inverted U-shaped functional relationship with brain MOR availability. Brain regions sensitive to daylength spanned the socio-emotional brain circuits, where MOR availability formed a spring-like peak. Causal effect of daylength on brain MOR availability was further verified by a *post hoc* experiment with repeated PET imaging of rats (n=9) under seasonal photoperiodic simulation. Therefore, the *in vivo* brain MOR availability in normal humans shows significant seasonal variation, which aligns with expected seasonal variation in mood and suicidality.

## Introduction

Seasonal rhythms profoundly impact mood. Negative affect including depression, anger, and hostility is at lowest during the summer ^1^, whereas seasonal affective disorder rates peak during the winter months ^2^. These changes are mediated by slow phasic changes in different neuroreceptor systems. For instance, long daylength increases brain serotonin and norepinephrine levels in mice and reduces depression and anxiety behaviour, compared to short daylength ^3^. Duration of daylight exposure is similarly associated with brain serotonin turnover in humans ^4^. The intimate link between seasonal fluctuations in mood and the contribution of opioidergic neurotransmission in human emotions ^5^ suggests potential seasonal variation of *in vivo* μ-opioid receptor (MOR) signalling.

Several lines of evidence suggest that MOR availability could vary seasonally in humans. Opioids are among the most commonly used illicit drugs in the US, where 2 % of the population have had opioid use disorder during their lifetime ^6^. Suicidal behaviour with prescription opioids follows a clear seasonal pattern, with suicide attempts peaking in the spring and fall ^7^. Further, post-mortem studies have established that suicide victims have increased MOR densities ^8,9^, and serious self-injurious behaviour is associated with increased plasma opioid growth factor levels ^10^. Although these studies have not directly assessed seasonal effects, it is well established that suicide rate peaks often in the spring regardless of the geographical location of the country ^11^. Finally, MORs are potent modulators of feeding ^12–14^ due to their contribution of hedonic or “liking” responses in the brain ^15^, and human feeding patterns show seasonal variation with caloric intake of fats typically peaking in the fall or winter ^16,17^. However, direct *in vivo* evidence on the causal seasonal effects on opioidergic neurotransmission is currently lacking.

Animal studies also suggest that there is seasonal variation in MOR-dependent signalling. The endogenous opioid signalling is crucial for the photoperiodic control of the seasonal reproductive cycle in mammals ^18^. Endogenous opioids inhibit the release of gonadotropin and sex hormones, and this inhibitory effect is enhanced under short daylength compared to long daylength ^19^. Also, MOR expression in hamster testes is increased during short days, with increased inhibitory control for testosterone secretion ^20^. In Siberian hamsters, brain expression of dynorphin A, an endogenous peptide agonist for opioid receptors, is increased under relatively longer daylength compared to shorter daylength ^21^, suggesting a potential impact of daylength on opioidergic signalling in the brain. Furthermore, effects of morphine on feeding behaviour of ground squirrels vary in accord to the hibernation state ^22^, indirectly reflecting a seasonal variation of MOR signalling.

Here, we tested whether seasonal variation in daylength, a noise-free cue for seasonal rhythms, influences MOR availability in the brain. MOR availability was quantified using *in vivo* positron emission tomography (PET) with MOR-sensitive agonist ligand [^11^C]carfentanil. We first conducted a cross-sectional study with previously acquired human PET imaging data to test whether there was seasonal difference in MOR availability. We then investigated experimentally whether seasonal variation in daylength causally influences brain MOR availability in rats. While rats underwent daylength cycle simulating seasonal changes, repeated [^11^C]carfentanil PET imaging was conducted (Supplementary Fig. 1). Our data suggest that there is seasonal variation in brain MOR availability in both humans and rats, with MOR availability peaking in seasons with intermediate daylengths.

## Results

### Daylength associated brain MOR availability in Humans

Figure 1A shows the mean MOR distribution in the human brain. We examined the association between natural variation in daylength (Fig. 1B) and MOR availability in humans. Brain full volume analysis revealed a statistically significant quadratic polynomial effects in a large brain cluster (39000 voxels; Fig. 1C-D), spanning the cingulate cortex, the superior frontal, medial frontal, middle temporal and superior temporal gyrus, insula and orbitofrontal cortex. Brain MOR availability in the brain cluster peaked at 15-19 hours (Fig. 1D).

**Figure 1.**
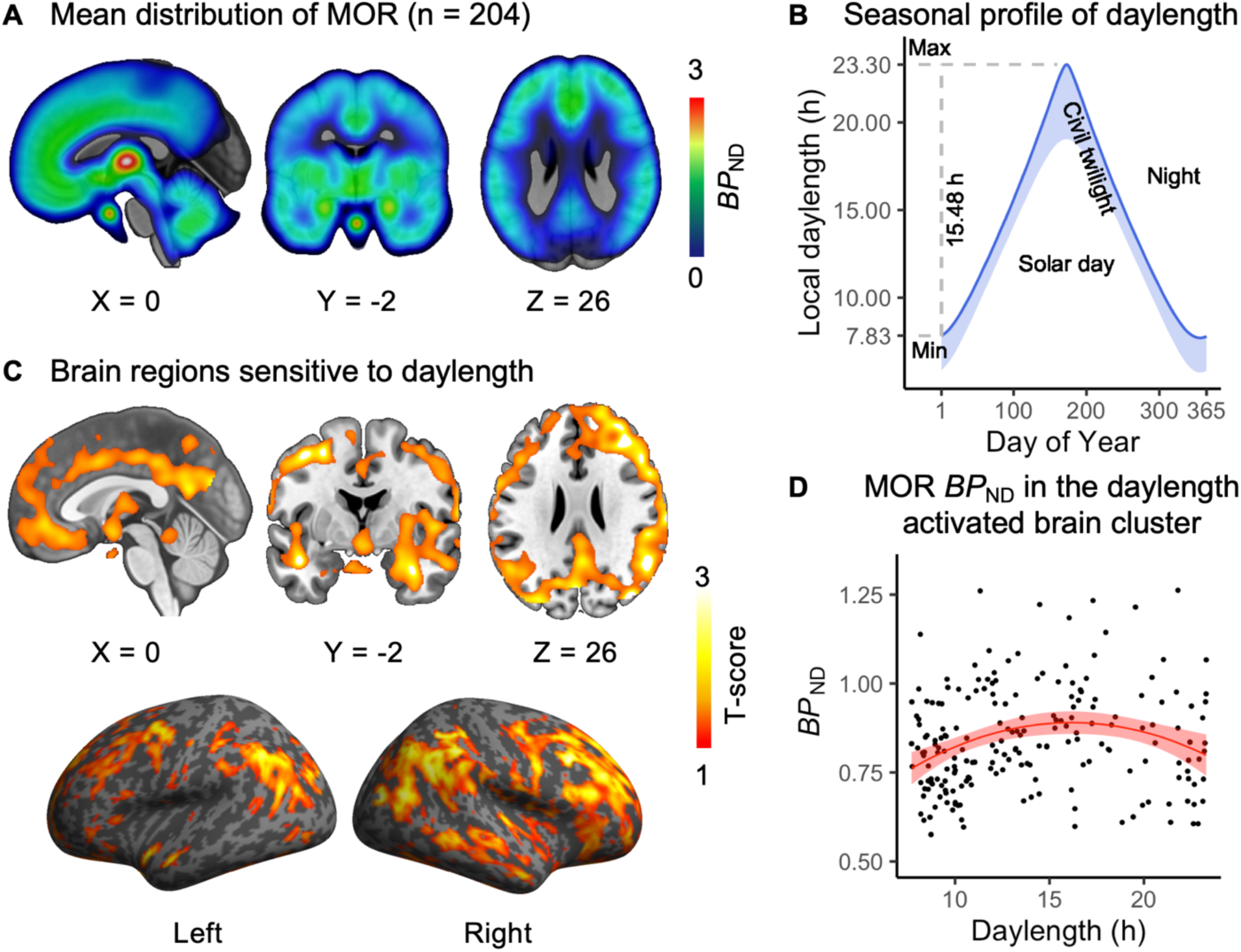
Human brain MOR availability and natural variation in daylength. **A.** Mean distribution of MOR availability in the subjects. **B.** Local seasonal distribution of daylength (day + civil twilight). **C.** The brain cluster sensitive to the seasonal daylength changes in quadratic polynomial regression model. p < 0.05, FDR-corrected. **D.** [^11^C]carfentanil *BP*_ND_ in the brain cluster as a function of daylength. Red line shows polynomial LS regression curve (y ∼ x + x^2) predicting *BP*_ND_, and shaded area shows 95% CI.

In addition, we did brain region of interest (ROI) analysis by involving thirty structurally determined brain ROIs (see Method). While involving all ROIs in the statistical analysis, both daylength (β = 0.044, 95% CI [0.008, 0.08]) and squared daylength (β = -0.0014, 95% CI [-0.0026, -0.00021]) were significant predictors for [^11^C]carfentanil *BP*_ND_, indicating an inverted-U shape functional relationship. No interaction effects between daylength and squared daylength with brain hemisphere (left versus right) or ROIs were found. Effect of daylength on individual ROIs is shown in Supplementary Table 1.

### Meteorological seasons and brain MOR availability in Humans

Traditionally defined seasons, according to the northern meteorological definition, including spring (March, April, May), summer (June, July, August), fall (September, October, November), and winter (December, January, February) were used as predictors for MOR *BP*_ND_ in the brain cluster (Fig. 1C-D). Each season is dummy-coded in a linear regression model, and we found that spring predicted higher brain MOR availability (β = 0.08, 95% CI [0.02, 1.14]), Fig. 2A. No other seasons had significant effects on MOR availability. In the local region, spring is with daylength of 16.39 ± 2.78 h (Fig. 2B). Therefore, MOR availability peaking at daylength 15-19 hours (Fig. 1D) suggests a spring-like peak.

**Figure 2.**
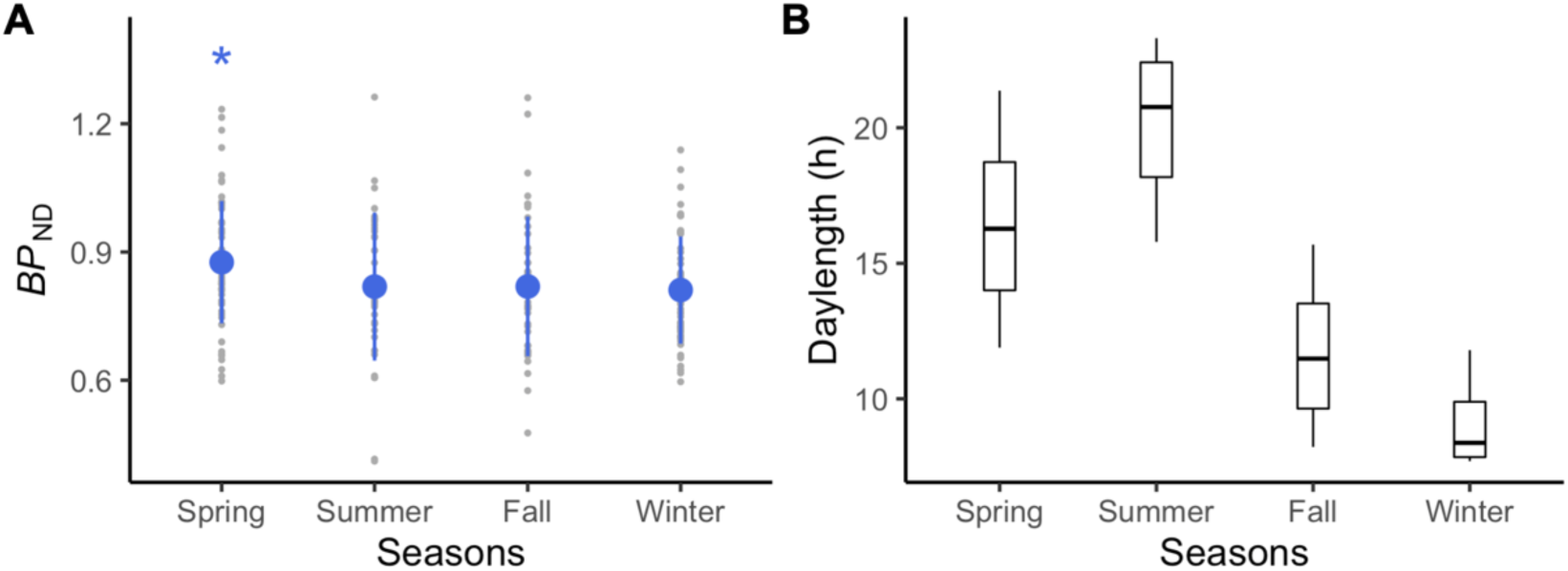
Spring is associated with highest brain MOR *BP*_ND_ in humans. **A.** MOR availability in the daylength activated brain cluster is plotted to seasons. Blue dots stand for means and error bars for standard deviations. **B.** Local daylength distribution in different seasons. * = p < 0.05.

### Causal effect of daylength on brain MOR availability in Rats

In addition to analysis of the human database, we set up a rat model PET imaging experiment to verify potential causal effect of daylength on brain MOR availability. This is also a crucial complement considering the quasi-experimental design nature of human data analysis which could not differentiate the effect of daylength from other highly correlated seasonal factors such as temperature. The rat model study consisted of experimental groups (n = 9) where daylength simulated the local daylength changes (no twilight) and control groups (n = 3) where daylength were kept consistent (12h light ON). When both experimental and control groups were included in the statistical model, both daylength (β = 0.16, 95% CI [0.035, 0.28]) and squared daylength (β = -0.0067, 95% CI [-0.012, -0.0017]) were significant predictors; the MOR availability and daylength had an inverted U-shaped relationship. Age, sex or group did not influence MOR availability, and no interaction effects were found. Analysis excluding the control group gave similar results (see Supplementary data).

We also modelled regional *BP*_ND_ separately for the experimental group rats (within-animal design), using fixed-effect factors daylength and squared daylength and varying intercepts for each rat. Daylength and squared daylength were significant predictors for [^11^C]carfentanil *BP*_ND_ in all regions (Fig 3; Supplementary Table 2). Including age and sex in the models did not improve the models, as evidenced by higher Akaike Information Criterion (AIC) values (Supplementary results & Supplementary Table 3). In addition, rats in the experimental versus control group showed slower weight gain (Supplementary Fig. 3) and had a higher level of blood corticosterone (β = 44.93, 95% CI [12.37, 77.48]).

**Figure 3.**
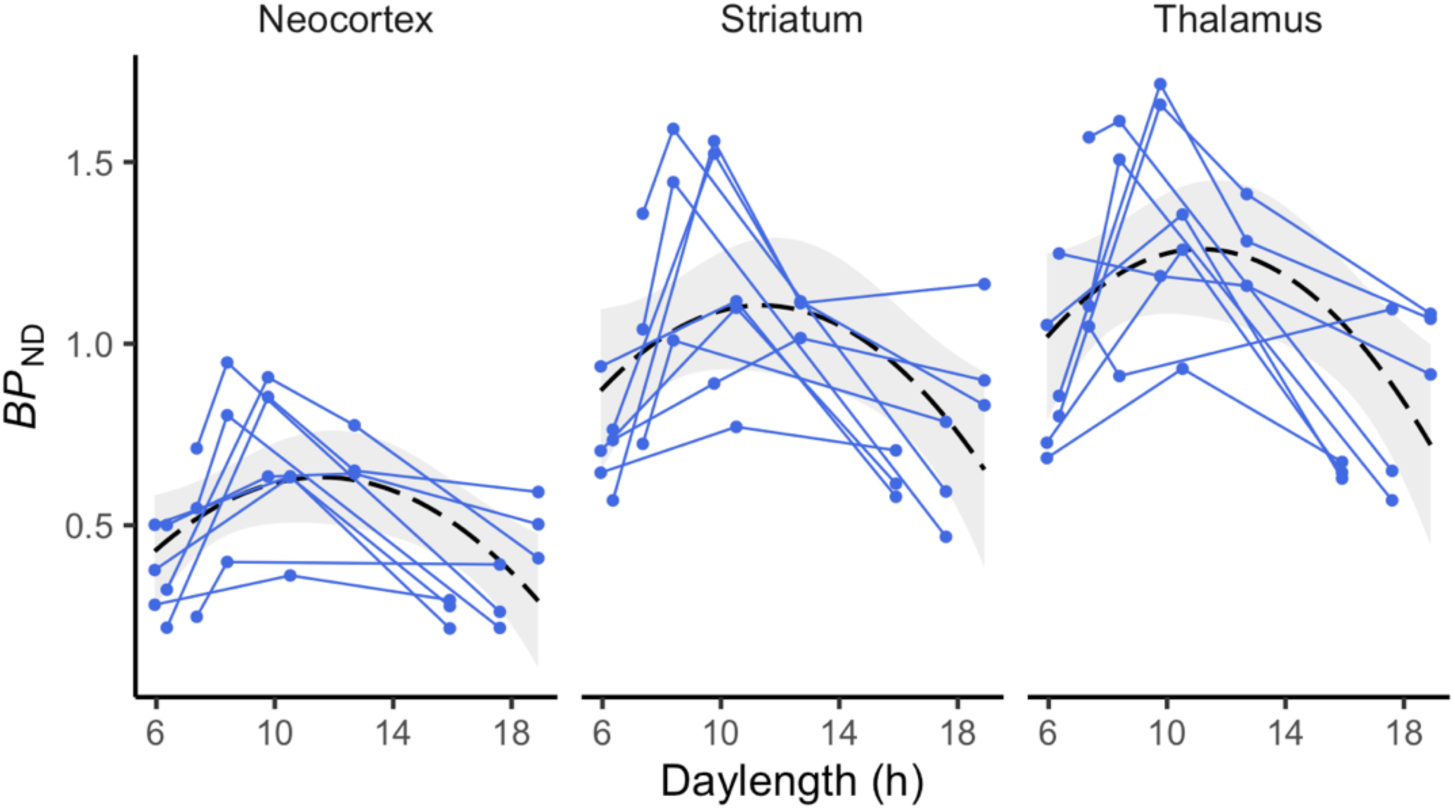
Within-animal changes in brain MOR *BP*_ND_ as a function of daylength. Individual animals are shown as separate lines. Black dashed line shows the LS-regression curve for the polynomial model (y ∼ x + x^2) predicting *BP*_ND_ across the whole sample. Shaded area shows 95% CI for the LS curve.

## Discussion

The current study reveals that seasonally varying daylength influences in vivo brain MOR availability in healthy humans. Increasing daylength modulated brain MOR availability following an inverted-U shape, with highest MOR bindings observed in intermediate daylengths. In rats, daylength causally influences MOR availability paralleling the quasi-experimental study in humans using a large database of historical PET imaging data. Most profound seasonal variation of MOR availability was found in large brain clusters including the amygdala, striatum, thalamus, cingulate cortex, orbitofrontal cortex, superior frontal and temporal cortex. Previous studies have established the role of seasonal variation in serotonin signalling on brain affective functions and pointed towards the role of daylight duration on its seasonal rhythm ^3,4^. The present data show that also MOR system undergoes significant seasonal variation, which might be linked with seasonal variation in mood and sociability.

The human data revealed salient seasonal variation of MOR availability in brain regions linked with emotional and social processing, such as amygdala, cingulate cortex and posterior superior temporal cortices ^5^. To our knowledge, this is the first *in vivo* evidence indicative of seasonal variation in MOR system in humans, and it fits with other types of evidence implying seasonal effects on MOR. In high-latitude regions daylight changes influence sleep quality, mood, and social behaviour ^23–26^. Prevalence of seasonal affective disorders or subsyndromes (time-locked to short daylength) in the local region is remarkably high ^27^. The spring-like peak of MOR availability (Fig. 2A) parallels with seasonal variation in opioid-modulated emotional changes such as mood ^1^, suicidality ^11^, particularly with prescription opioids ^7^ that peak particularly in the spring. Considering the impact of photoperiod on seasonal affective disorders ^28^ as well as the intimate linkage between MOR signalling and social and affective functions ^5,29^, the current study suggests that seasonal variation in MOR availability could be a potential brain mechanism under seasonal affective changes.

The rat experiment finding demonstrates that daylength causally influences brain MOR availability, verifying the human data findings, and goes in line with previously reported impact of photoperiod on MOR dependent signalling in mammals ^19–22^. Exact mechanism on the daylength-mediated alterations in brain MORs is however unclear. In mammals, neural responses to daylength are largely mediated by the superchiasmatic nucleus (SCN). The SCN generates endogenous neural signal, and adjusts the duration of this signal in accord to changes of the solar day length as is detected via the retinohypothalamic tract ^30,31^. While MORs are most widely expressed in the subcortical and limbic brain regions, it is possible that daylength affects MOR signalling via the retinohypothalamic tract, calling for in-depth studies.

In rats, we also found that seasonally alternating daylength slowed weight gain and increased blood corticosterone levels in adult rats (in comparison with rats kept in a fixed 12-hour light cycle), in line with previous findings ^32,33^. Daylength-dependent changes in corticosterone levels may be due to elevated stress caused by extremely long daylength, as supported by a positive effect of daylength on blood corticosterone levels in rats (Supplementary results). In humans, exposure to extremely long daylength in summer also increases human blood cortisol levels ^34^. Blood corticosterone levels have been linked with social stress and activity of MOR signalling in mammals ^35^, where increased MOR avidity is along with lower cortisol level under stress. This is similarly supported by our finding that brain MOR availability in rats is negatively associated with blood corticosterone levels (Supplementary Fig. 4).

While the current study cannot disentangle the relation between corticosterone levels and MOR signalling in humans, the findings suggest that social and physiological stress could be linked with MOR availability. Regardless of the euphoric effect of longer daylength ^1,28^, cerebral MOR availability did not increase linearly with daylength but demonstrated an inverted-U shape, potentially due to the increased physiological or psychological stress under extremely long daylength. This link between MOR signalling and stress is corroborated by the fact that opioid agonists and partial agonist buprenorphine reduce symptoms of anxiety and depression in rats ^36^ as well as alleviate the effects of psychological stressors such as separation distress ^37^. In line with this, human molecular and functional imaging studies have found that high opioidergic tone suppresses hemodynamic brain responses to viewing distressing videos ^38,39^.

While PET being the optimal non-invasive approach to quantify *in vivo* brain MOR availability, drawbacks exist in the current study. The human study was based on historical data, where each subject was imaged only once, and quasi-experimental design (natural changes in daylight), as longitudinal multi-scan studies would yield a significant radiation load. Contribution of other seasonal factors (such as temperature) cannot however be ruled out in the human data, whereas we verified the causal effect of daylength on brain MOR availability using animal experiments. Furthermore, daylength was estimated using astronomical data without considering day-to-day (e.g. cloud cover) and individual variation (e.g. time spent outside) in light exposure. Nevertheless, we had a large sample and the results paralleled those from the experiment done in rats, albeit with smaller effect size. The human data were also sampled from different projects and scanners, which were corrected for potential scanner-related biases in the analyses. Technically, while [^11^C]carfentanil *BP*_ND_ in baseline condition is proportional to MOR density, the exact contributions of MOR density, receptor affinity, and baseline occupancy by endogenous opioids cannot be assessed in a single measurement ^40,41^, and these components cannot be differentiated in a single scan. [^11^C]carfentanil is an agonist tracer preferably binding MORs in the high-affinity state ^41^ and has low test-retest variability in PET imaging ^42^. Although endogenous opioids compete for binding sites with [^11^C]carfentanil ^43,44^, the basal opioid concentrations are often very low at least in rats ^45^. Accordingly, [^11^C]carfentanil *BP*_ND_ in baseline likely reflects density of MORs. Finally, current findings based on human data should be interpreted while keeping in mind the large magnitude of local photoperiodic variation, which should be cautiously interpolated to regions with lower latitudes.

Taken together, we conclude that seasonal variation in daylength influences brain MOR availability, following an inverted U-shaped curve and peaking at 15-19 hours in humans. Given the intimate links between MOR signalling and socioemotional behaviour ^29,46^, these results suggest that the MOR system might underlie seasonal variation in human mood and social behaviour ^1,2^, and imply that MOR might be a feasible target for treating seasonal affective disorders.

## Methods

### Study with human PET image database

#### Subjects

Human study was performed using retrospective analysis of historical subjects scanned with PET using radioligand [^11^C]carfentanil at different times of the year (Supplementary Fig. 1). The data were retrieved from the AIVO database (http://aivo.utu.fi) of *in vivo* molecular images hosted by Turku PET Centre. We identified all the baseline scans from individuals with no neurologic or psychiatric disorders who had been scanned between 2003 and 2018. The resulting data frame of 204 individuals (132 males, 72 females; age 32.4 ± 10.8 y) consists of scans from 11 research projects and five PET scanners, with details of the subjects described in ^47^. No subjects abused alcohol or illicit drugs or used medications affecting the central nervous system.

#### PET image analysis

PET data were analysed and modelled using Magia toolbox (https://github.com/tkkarjal/magia) ^48^, an automated processing pipeline developed at the Turku PET Centre running on MATLAB (The MathWorks, Inc., Natick, Massachusetts, United States). Pre-processing consisted of framewise realignment and co-registration of the PET and magnetic resonance images. Tracer binding was quantified using *BP*_ND_ ^49^, which is the ratio of specific binding to non-displaceable binding in tissue. *BP*_ND_ was estimated using simplified reference tissue model with occipital cortex as the reference region ^50,51^. Parametric *BP*_ND_ images were also calculated for full-volume analysis. They were spatially normalized to MNI-space via segmentation of T1-weighted MRIs and smoothed with an 8-mm Gaussian kernel. Regions of interest, including the reference region, were parcellated for each subject using FreeSurfer (http://surfer.nmr.mgh.harvard.edu/).

#### Full-volume human data analysis

For each subject, daylength was calculated as the daytime plus civil twilight on the day when the PET image was acquired. Civil twilight involves morning civil twilight that begins when the geometric centre of the sun is 6° below the horizon and ends at sunrise and evening civil twilight that begins at sunset and ends when the geometric centre of the sun reaches 6° below the horizon. Calculation was done using R package “suncalc”, where calculations were based on geographic location of the Turku PET Centre (Turku, Finland; latitude = 60.4518, longitude = -22.2666) and formulas provided by Astronomy Answers articles about position of the sun and the planets (https://www.aa.quae.nl/en/reken/zonpositie.html).

We first modelled the effect of daylength on MOR availability using multiple regression, as implemented in SPM12 (http://www.fil.ion.ucl.ac.uk/spm/). Polynomial expansion of daylength to the second order and linear component of daylength were used as regressors. Subject age, sex, scanner and body mass index (BMI) were used as nuisance covariates. MOR *BP*_ND_ values in the daylength-sensitive brain cluster were also estimated using the MarsBaR toolbox ^52^, and plotted as a function of daylength and seasons for visualization.

#### Region of interest analysis

We also analysed the regional MOR availability in 15 brain ROIs (based on subject-wise FreeSurfer delineations) for both hemispheres (in total 30 ROIs), as listed in Supplementary Table 1. Pooled ROI values were analysed using linear mixed-effects model with varying intercepts for each subject. Fixed effect factors included daylength, squared daylength, age, sex, BMI, scanner type, and their interaction effects with ROI and brain hemisphere (right vs. right). In the single ROI data analysis, linear regression model was used with factors including daylength, squared daylength, age, sex, BMI, and scanner type. Regional MOR *BP*_ND_ was log-transformed in the statistical analysis as previously ^47^. R statistical software (version 3.6.0) using lme4 package were used in analysing the human ROI data.

### Rat model PET imaging study

#### Animal handling and seasonal simulation

The National Animal Experiment Board of Finland approved all the procedures and protocols (license number ESAVI/8355/2019) in accordance with the EU Directive 2010/63/EU on the protection of animals used for scientific purposes. Eighteen adult (age > 90 days) Sprague-Dawley rats (twelve of them were PET imaged as described in the next section; 11 males, 7 females; Central Animal Laboratory, University of Turku, Turku, Finland) were housed under controlled laboratory conditions in open top cages with free access to CRM-E diet (SDS, UK) and water. Rats were caged in groups of two or three same-sex rats. No rats were removed before finishing the last PET imaging to keep the social environment stable and to avoid additional stress ^53^. For the rats in the experimental group (9 males, 5 females), in-house light with variable ON/OFF duration was programmed, simulating the local seasonal change of daylength with a speeded cycle completed in three months (Supplementary Fig. 1). Rats in the control group (2 males, 2 females) were kept in a different room with constant daylength cycle (12h ON/ 12h OFF, no twilight), with all other conditions the same with the experimental group. The same type of LED lighting was used for control groups and the experimental groups.

#### PET imaging and processing

Twelve out of eighteen rats (experimental group: 6 males, 3 females; control group: 2 males, 1 female) were studied with dynamic [^11^C]carfentanil PET imaging for 3-4 times under isoflurane anaesthesia. Radiotracers were divided by three rats at each scanning day (2 males, 1 female), and via using two different scanners to maximize the data collection. Due to larger body size, Inveon Multimodality PET/CT (Siemens Medical Solutions, Knoxville, TN, USA) was used for imaging the male rats (two rats each time), whereas Molecubes PET/CT (Molecubes, Gent, Belgium) was used for the female rats. In total, there were 42 PET/CT scans. Rats were weighed on the scanning day. For the Inveon Multimodality PET/CT scanner, with relatively lower resolution and sensitivity, the aimed dosage of radiotracer was 5 MBq, while for Molecubes PET/CT scanner rats were scanned with aimed dosage of 1 MBq. Accordingly, sex effects cannot be separated from scanner type and dosage dependent effects.

Dynamic PET images were analysed using Carimas software (version 2.10.3.0) developed at the Turku PET Centre, Finland. The PET data sets were reconstructed into 20 time frames using OSEM3D algorithm: 6 × 0.5 minutes, 3 × 1 minutes, 4 × 3 minutes, and 7 × 6 minutes. We first created a template of structural CT image and brain atlas. In the template, a rat’s skull CT image was aligned with the Waxholm Space Atlas of the Sprague Dawley rat brain ^54^ to be used as the reference for the normalization of the brain scans. We first visually checked that the PET and CT images were in alignment. Next, PET and CT images were locked regarding their positions using Carimas. Individual skull images were then manually realigned and resized to the template CT image. Once the skull image was scaled with the CT image, the CT and the PET images were rigidly co-registered with the CT template (Supplementary Fig. 2).

Simplified reference tissue model was used for estimating of non-displaceable MOR binding potential (*BP*_ND_) in the neocortex, striatum and thalamus ^49^. Cerebellum was used as the reference region because it is devoid of MOR in rats (see Supplementary Fig. 5 for *ex vivo* biodistribution results). We excluded the first five minutes from model fitting to rule out confounding perfusion effects. Time-activity curves (TACs) from brain regions of interest (thalamus, striatum and neocortex) defined by the Waxholm atlas were then extracted for data analysis.

#### Serum sampling and measurement of corticosterone levels

Blood samples were collected by lateral tail vein puncture at 9:00-11:00 am. For experimental group rats, samples included collections conducted one day prior to the PET imaging (30 samples out of 33). For the control group, sample collections were randomly conducted. Altogether, 51 blood samples were obtained (18 for the control group). The samples were centrifuged at 7000 rpm for 1.5 minutes and the serum was stored at -20°C. Levels of serum corticosterone were measured using an ELISA kit (Enzo Life Sciences, Ann Arbor, MI, USA).

#### Statistical analysis

Data were analysed with R (version 3.6.0) using lme4 package ^55^. To estimate the effects of daylength on MOR availability, we used a linear mixed-effects model with varying intercepts for each rat. Fixed effect factors included daylength, squared daylength, age, sex, group, and the interaction effects between regions of interests (ROI) and all fixed effect factors were included.

## Author contribution

LS, JT, EY, AR designed the rat imaging experiment. LS analysed the data under supervision of JT, AR, and LN. All authors wrote the manuscript. HL, AH, JV, SH conducted the rat imaging experiments under supervision of EY and AR. KE and ES measured the corticosterone levels. JI, TKarjalainen, TKantonen, PN, JH, VK, KK, JH, HS, JOR, LN contributed to construction of the human brain database. All the authors discussed the results and commented on the manuscript.

## Acknowledgements

The study was supported by the Academy of Finland (grant #317680 to JT, #294897 and #304385 to LN, and #310962 to JOR), Sigrid Juselius Foundation, and Jane and Aatos Erkko Foundation (to AR). LS is supported by Turku Collegium for Science and Medicine, University of Turku. TKantonen is supported by Finnish Cultural Foundation (Southwest Finland Fund) and Emil Aaltonen Foundation. JOR is supported also by the Sigrid Juselius Foundation and Finnish State Research Funding. We thank Sauli Piirola, for help with Carimas scripting, and Elina Kahra for her assistance in laboratory analysis of blood corticosterone levels. We thank Paulina Chrusciel, Ella Kujala, Malgorzata Major and Joonas Khabbal from UTUCAL for assisting lighting planning and blood sampling.

