## supplements for "Seasonal variation in brain mu-opioid receptor availability"

### Supplementary methods

#### Study design

The current study consists of human brain PET image database analysis and an experimental rat model study (Supplementary Fig. 1).

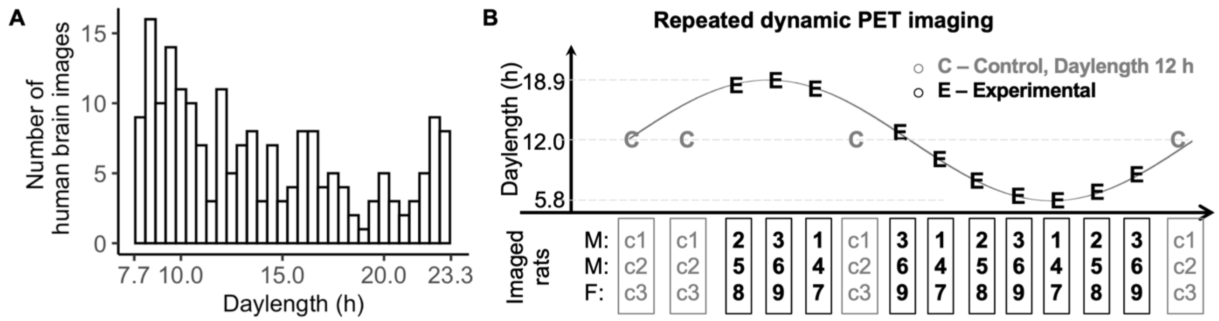


**Supplementary Figure 1**. Study design. **A**. Number of human PET scans at different daylengths (1 bin = 0.5 h). **B**. Experimental design for the rat model PET study. M = males, F = females, C = control

#### Rat model study

*Ex vivo gamma counting*

Twenty minutes after the injection of [$^{11}$C]carfentanil (40 MBq), rats were sacrificed under isoflurane anaesthesia and samples of various tissues and brain regions were excised, weighed, and measured for radioactivity using a gamma counter (Triathler 3", Hidex, Turku, Finland). The results are shown as percentage of injected radioactivity dose per gram of tissue (%ID/g).

*PET image processing*

Rat PET brain images were pre-processed as shown in Supplementary Fig. 2.

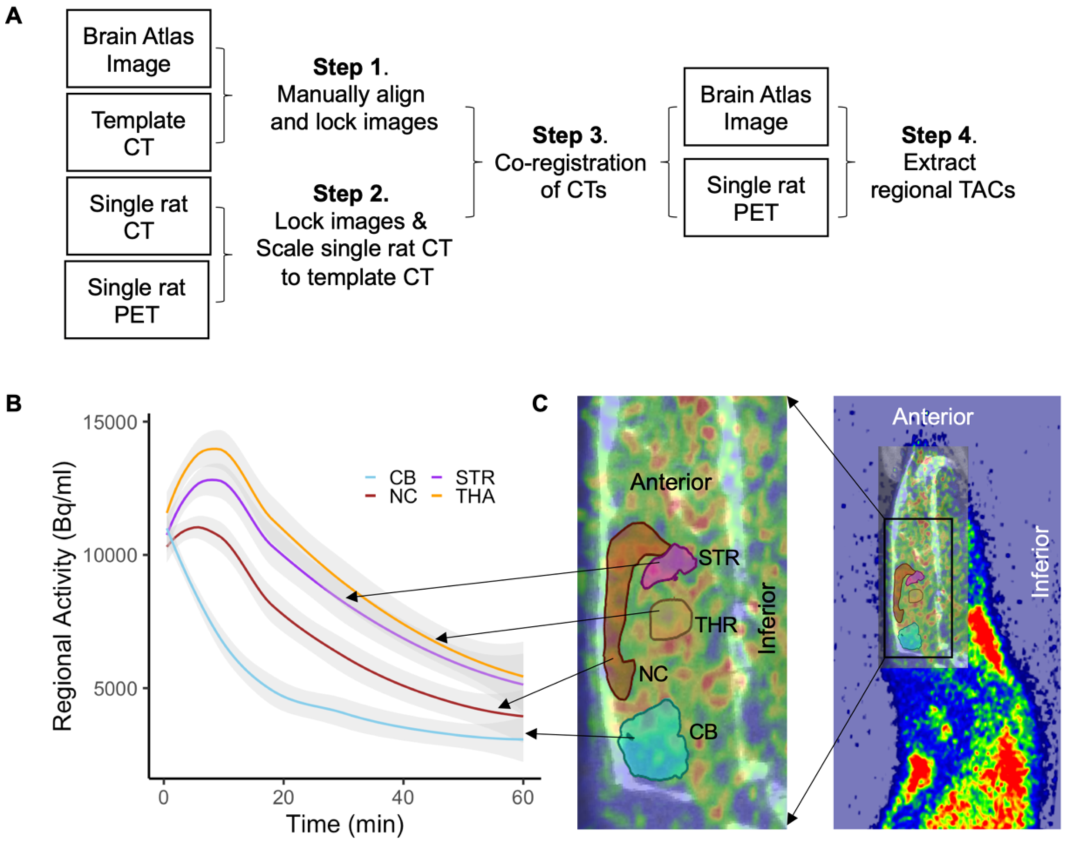


**Supplementary Figure 2**. Image processing and analysis for rat *in vivo* PET data. **A.** Schematic description of the image processing procedures. **B.** Mean (with shaded area for 95% CI) time activity curves for the reference region and the target regions extracted using (**C**) ROI template image overlaid on the PET and CT images. CB = cerebellum, STR = striatum, NC = neocortex, THA = thalamus.

### Supplementary Results

#### Findings in human database analysis

*Human brain ROI analysis*

Effect of daylength on MOR $BP_{ND}$ on regional binding was done for each ROI, using factors daylength, squared daylength, age, sex, scanner type and BMI. Both daylength and squared daylength were significant predictors for most of the ROIs. Uncorrected results for each ROI are listed in Supplementary Table 1.

**Supplementary Table 1.** Effect of daylength and squared daylength on regional MOR binding in humans (uncorrected for multiple comparison). Significance levels of p values are marked: * = "p<0.05", ** = "p < 0.01", *** = "p < 0.001". Effect of age and adjusted $R^2$ of the full models are also listed. ACC = anterior cingulate cortex, MTP = mid-temporal gyrus, PCC = posterior cingulate cortex, SFG = superior frontal gyrus.

| Brain Hemisphere | Region | Daylength | | Squared daylength | | Age | | adjusted $R^2$ |
|---|---|---|---|---|---|---|---|---|
| | | β | 95% CI | β | 95% CI | β | 95% CI | |
| right | amygdala | 0.03 | [-0.005, 0.07] | -0.001 | [-0.002, 0.0001] | -0.005 | [-0.008, -0.002] | 0.20 |
| right | caudate | 0.03 | [-0.01, 0.07] | -0.0009 | [-0.002, 0.0004] | 0.004 | [0.0002, 0.007]* | 0.28 |
| right | dorsal ACC | 0.03 | [-0.001, 0.07] | -0.001 | [-0.002, -0.00005]* | 0.003 | [-0.0001, 0.006] | 0.32 |
| right | inferior MTG | 0.04 | [-0.003, 0.08] | -0.001 | [-0.002, 0.0002] | 0.01 | [0.007, 0.013]*** | 0.33 |
| right | insula | 0.04 | [0.001, 0.07]* | -0.001 | [-0.002, -0.00002]* | 0.002 | [-0.0007, 0.005] | 0.09 |
| right | middle MTG | 0.05 | [0.01, 0.09]** | -0.002 | [-0.003, -0.0004]* | 0.009 | [0.007, 0.01]*** | 0.34 |
| right | nucleus accumbens | 0.03 | [-0.008, 0.06] | -0.0009 | [-0.002, 0.0002] | -0.002 | [-0.005, 0.0005] | 0.11 |
| right | orbitofrontal cortex | 0.04 | [0.005, 0.08]* | -0.001 | [-0.002, -0.0001]* | 0.007 | [0.004, 0.01]*** | 0.28 |
| right | pars opercularis | 0.05 | [0.02, 0.09]** | -0.002 | [-0.003, -0.0005]** | 0.006 | [0.003, 0.009]*** | 0.31 |
| right | PCC | 0.04 | [0.04, 0.08]* | -0.001 | [-0.002, -0.0001]* | 0.004 | [0.001, 0.007]** | 0.34 |
| right | putamen | 0.04 | [0.008, 0.08]* | -0.001 | [-0.003, -0.0003]* | 0.002 | [-0.0004, 0.005] | 0.17 |
| right | rostral acc | 0.04 | [0.007, 0.08]* | -0.001 | [-0.003, -0.0002]* | 0.005 | [0.002, 0.008]** | 0.26 |
| right | SFG | 0.04 | [0.006, 0.08]* | -0.001 | [-0.003, -0.0002]* | 0.006 | [0.002, 0.009]*** | 0.38 |
| right | temporal pole | 0.05 | [0.01, 0.1]* | -0.002 | [-0.003, -0.0003]* | 0.006 | [0.002, 0.01]** | 0.16 |
| right | thalamus | 0.03 | [0.00003, 0.06]* | -0.001 | [-0.002, -0.00005]* | -0.005 | [-0.008, -0.003]*** | 0.19 |
| left | amygdala | 0.04 | [0.008, 0.08]* | -0.001 | [-0.003, -0.0002]* | -0.008 | [-0.01, -0.005]*** | 0.28 |
| left | caudate | 0.03 | [-0.01, 0.07] | -0.0009 | [-0.002, 0.0005] | 0.003 | [-0.00007, 0.007] | 0.26 |
| left | dorsal ACC | 0.03 | [-0.007, 0.06] | -0.001 | [-0.002, 0.0002] | 0.002 | [-0.0009, 0.005] | 0.31 |
| left | inferior MTG | 0.04 | [-0.002, 0.07] | -0.001 | [-0.002, 0.0001] | 0.01 | [0.007, 0.013]*** | 0.31 |
| left | insula | 0.03 | [-0.008, 0.06] | -0.0009 | [-0.002, 0.0002] | 0.002 | [-0.0004, 0.005] | 0.16 |
| left | middle MTG | 0.03 | [-0.008, 0.07] | -0.0009 | [-0.002, 0.0003] | 0.01 | [0.008, 0.014]*** | 0.16 |

| | | | | | | | | |
|---|---|---|---|---|---|---|---|---|
| left | nucleus accumbens | 0.03 | [-0.01, 0.06] | -0.0008 | [-0.002, 0.0004] | -0.002 | [-0.005, 0.0006] | 0.11 |
| left | orbitofrontal cortex | 0.05 | [0.008, 0.08]* | -0.001 | [-0.003, -0.0002]* | 0.008 | [0.005, 0.01]*** | 0.32 |
| left | pars opercularis | 0.03 | [-0.007, 0.07] | -0.001 | [-0.002, 0.0002] | 0.006 | [0.003, 0.009]*** | 0.36 |
| left | PCC | 0.05 | [0.01, 0.08]** | -0.002 | [-0.003, -0.0004]** | 0.005 | [0.002, 0.008]*** | 0.37 |
| left | putamen | 0.03 | [-0.002, 0.06] | -0.001 | [-0.002, 0.00008] | 0.003 | [0.0029, 0.005]* | 0.18 |
| left | rostral acc | 0.03 | [-0.003, 0.07] | -0.001 | [-0.002, 0.0001] | 0.005 | [0.003, 0.008]*** | 0.27 |
| left | SFG | 0.04 | [0.002, 0.08]* | -0.001 | [-0.003, -0.00005]* | 0.006 | [0.003, 0.01]*** | 0.40 |
| left | temporal pole | 0.03 | [-0.01, 0.07] | -0.001 | [-0.002, 0.0004] | 0.008 | [0.004, 0.01]*** | 0.16 |
| left | thalamus | 0.03 | [-0.0004, 0.06] | -0.001 | [-0.002, -0.00001]* | -0.006 | [-0.008, -0.003]*** | 0.16 |

### Findings in Rat Model Study

*Effect of daylength on brain MOR $BP_{ND}$*

When including only the experimental group (without control group with constant daylength at 12h), daylength (β = 0.16, 95% CI [0.029, 0.28]) and squared daylength (β = -0.0058, 95% CI [-0.011, -0.00057]) had significant effect on MOR binding, generating an inverted-U shape. Age, sex or group did not influence MOR binding and no interaction effects were found.

Regional $BP_{ND}$ for the experimental group rats were separately modelled using fixed-effect factors daylength and squared daylength, Supplementary Table 2.

**Supplementary Table 2.** Daylength and squared daylength as predictors for brain regional MOR binding in rats.

| ROI | Factor | β value | 95% CI | t score | p value | Marginal $R^2$ |
|---|---|---|---|---|---|---|
| Neocortex | Daylength | 0.2 | [0.1, 0.28] | 4.14 | 0.002 | 0.33 |
| | Squared daylength | -0.008 | [-0.012, -0.0045] | -4.27 | 0.002 | |
| Striatum | Daylength | 0.25 | [0.11, 0.38] | 3.57 | 0.002 | 0.26 |
| | Squared daylength | -0.01 | [-0.017, -0.005] | -3.64 | 0.002 | |
| Thalamus | Daylength | 0.25 | [0.11, 0.4] | 3.43 | 0.005 | 0.33 |
| | Squared daylength | -0.01 | [-0.017, -0.005] | -3.75 | 0.003 | |

ROI-wise models were also run with full model including daylength, squared daylength, age and sex as fixed-effect factors. Both daylength and squared daylength were significant predictors in all ROIs, Supplementary Table 2. These regional models were compared with the model using only daylength and squared daylength as predictors using Akaike information criterion, and in all regions the full model, in comparison yielded higher AIC values (Neocortex: 38.66 versus 24.28; Striatum: 60.17 versus 48.16: Thalamus: 61.30 versus 48.16).

**Supplementary Table 3.** Results of ROI analysis using full models including age and sex as factors.

| ROI | Factor | beta | 95% CI | t score | p value | Marginal $R^2$ |
|---|---|---|---|---|---|---|
| Neocortex | Daylength | 0.20 | [0.12, 0.29] | 4.61 | 0.001 | 0.39 |
| Neocortex | Squared daylength | -0.008 | [-0.012, -0.0047] | -4.46 | 0.0005 | |
| Striatum | Daylength | 0.25 | [0.11, 0.39] | 3.49 | 0.007 | 0.30 |
| Striatum | Squared daylength | -0.01 | [-0.017, -0.004] | -3.64 | 0.008 | |
| Thalamus | Daylength | 0.28 | [0.14, 0.43] | 3.89 | 0.07 | 0.39 |
| Thalamus | Squared daylength | -0.01 | [-0.018, -0.006] | -3.79 | 0.12 | |

*Impact of seasonal cycle on weight gain*

To test whether the variable seasonal rhythm influenced weight gain, weight was analysed using fixed effect factors including age, sex and group, and using rats as the random effect factor. To see the effect of group on the rate of weight gain, we also included an interaction effect between group and age. We found that weight increases by aging (β = 0.96, CI [0.78, 1.15]), and males have higher weights (β = 198, 95% CI [117, 216]). There was also an interaction between age and group (β = -0.31, 95% CI [-0.58, -0.03]), suggesting that simulated seasonal changes in daylength influenced the growth rate of rats, Supplementary Fig. 3.

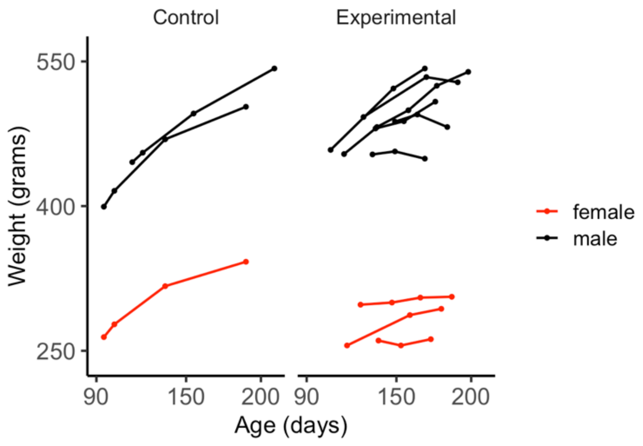


**Supplementary Figure 3.** Weight gain of rats under different conditions.

*Effect of experimental condition on blood corticosterone levels*

Serum corticosterone levels were analysed using fixed effect factors including daylength, age, sex and group, with rats as random factor. In addition to a group-level difference (see main text), daylength had positive effect on corticosterone levels (β = 7.6, 95% CI [2.22, 12.98]). Aging also led to increased blood corticosterone levels (β = 1.55, 95% CI [0.85, 2.25]) and male rats had lower corticosterone levels (β = -39, 95% CI [-71.43, -6.56]).

*Association between blood corticosterone and MOR $BP_{ND}$*

For experimental group rats, data of blood corticosterone levels include samples taken one day before the imaging date (not for the control group). MOR binding in brain regions of interest was modelled using varying intercept for rats and corticosterone level as the only fixed-effect factor. Corticosterone level was associated with MOR binding in the striatum (95% CI [-0.0033, -0.000078]), but not in neocortex (95% CI [-0.0022, 0.00025]) and thalamus (95% CI [-0.003, 0.00027]), Supplementary Fig. 4. However, the effect in striatum was not significant after Bonferroni correction for multiple comparison.

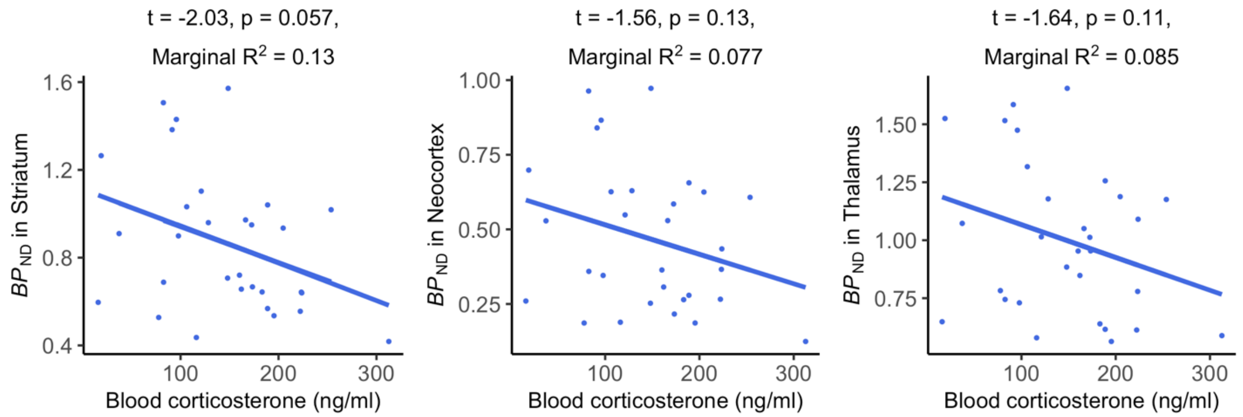


**Supplementary Figure 4.** Association between blood corticosterone levels and MOR $BP_{ND}$ in rats. Dot-plots represent association between blood corticosterone level and brain regional MOR $BP_{ND}$. Line plots (mostly overlapping) were obtained at individual levels for each ROI. Statistical indices on top of each plot illustrate results of the mixed-effect linear models to predict reverse MOR $BP_{ND}$ using corticosterone level as the only fixed-effect factor.

*Ex vivo gamma counting*

*Ex vivo* biodistribution was conducted for measuring MOR binding in different brain regions. This confirmed that cerebellum is a suitable ideal reference region for SRTM with [$^{11}$C]carfentanil (Supplementary Fig. 5)

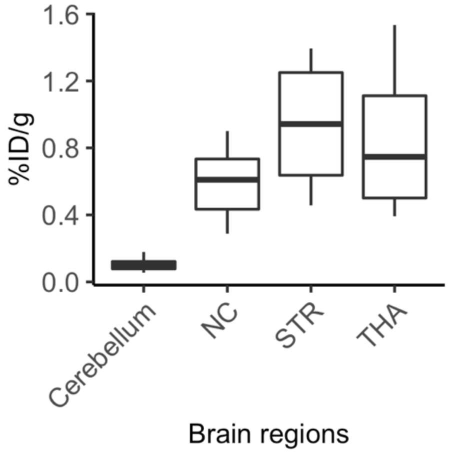


**Supplementary Figure 5.** *Ex vivo* biodistribution of [$^{11}$C]carfentanil in brain regions of interests.
